# Ontogeny of familiarity with foraging landscape and foraging abilities in the tropical social wasp *Ropalidia marginata*

**DOI:** 10.1101/272831

**Authors:** Souvik Mandal, Anindita Brahma, Raghavendra Gadagkar

**Author notes:** Correspondence: Souvik Mandal, Postal address: Centre for Ecological Sciences, Indian Institute of Science, Bangalore 560012, 12 India.

## Abstract

Possessing spatial familiarity with their foraging landscape may enable animals to reduce foraging effort without compromising on foraging benefits. For animals inhabiting feature-rich landscapes, spatial familiarity can increase with increasing age/experience. To check whether this holds for individually foraging tropical social wasp *Ropalidia marginata*, we recorded the number and duration of all foraging trips, the identity of the materials brought to the nest (building material, water or food) and the directions of outbound and inbound flights (respective to their nests) of known-age foragers from three natural colonies, each for three consecutive days. The average trip duration and time spent daily in foraging increased rapidly until about first four weeks of their life, during which they rarely brought food to their nest, although many of them brought building material and water. Thereafter, their average as well as per day duration of foraging trip started decreasing gradually. Nevertheless, their foraging efficiency and success for food kept on increasing monotonically with age. These results suggest that older wasps were more efficient in foraging despite spending less time doing so. With increasing age, wasps developed individual preferences for the direction of their outbound flights, increased directionality of their inbound trips as well as the angular difference between their outbound and subsequent inbound flights, indicating development of spatial memory. We conclude that wasps acquire familiarity with their foraging landscape in their initial foraging phase and gradually develop robust memory for rewarding sites and routes to those sites, which enables them to increase their foraging capabilities.

**SUMMARY STATEMENT:** Contrary to insects inhabiting less-featured landscapes, tropical social wasps invest weeks to get familiar with foraging landscapes during their early foraging lives. This eventually enables them to increase foraging gain with reduced effort.

## INTRODUCTION

Minimizing foraging effort without compromising foraging benefit is a perpetual challenge for foraging animals. For animals inhabiting feature-rich landscapes, learning and memorising features of foraging landscapes, especially those related to the locations of rewards may eventually enable them to collect substantial foraged materials with a minimal investment in foraging effort (Caraco, 1980; Kamil and Roitblat, 1985). It has been postulated that more the time an animal travels within a landscape, more would be its spatial familiarity with the surrounding (Boyer and Walsh, 2010), and thus, better would be its homing capabilities (Dyer, 1996). This in turn may enable them to attain higher foraging benefits (Bracis et al., 2015; Dukas and Visscher, 1994; Pyke et al., 1977; Raine and Chittka, 2008). On the other hand, foraging is a costly affair in terms of time, energy and associated risks like predation. Thus, it would be a proficient strategy for animals to strike an optimum balance between the time they spend to gain familiarity with their foraging landscapes and on foraging, and the amount of foraged materials (Abrams, 1991; Norberg, 1977).

To acquire such spatial familiarity, foragers of social hymenopterans like ants (Fleischmann et al., 2016), honey bees (Capaldi et al., 2000; Degen et al., 2015), and bumble bees (Osborne et al., 2013; Woodgate et al., 2016) begin their foraging life with a few exploratory flights/walks, and very soon they begin to forage for food. The Saharan desert ants, which scavenge in featureless and non-patchy desert landscapes, increase their foraging success as well as their foraging duration with the advancement of their short foraging life (Wehner et al., 2004). Similarly, honey bees inhabiting temperate landscape also increase foraging gain with age but they also cover more foraging area per trip and increase their flight speed (Capaldi et al., 2000). Thus, their foraging success seems to depend much on their current foraging effort (and not on the effort they put on the exploratory/learning phase), which might be an ideal strategy for foragers evolved in such less-featured non-patchy landscapes. On the other hand, landscapes with high density of features, like the tropics, probably offer much more spatial information but much less visual continuity (Cartwright and Collett, 1987; Zeil, 2012). Thus, insects inhabiting complex feature-rich terrains encounter homing challenges different from the insects inhabiting less-featured terrains „ and thus they may evolve with different homing and foraging strategies. Interestingly, whereas insects inhabiting less-featured landscapes rely heavily on their path integration system for homing, foragers of the Australian ant *Melophorus bagoti* (Narendra, 2007) inhabiting semi-desert cluttered landscape or the tropical ant *Gigantiops destructor* (Macquart et al., 2006) inhabiting complex tropical rain forests rely heavily on the learnt visual features of their foraging landscape. However, we are yet to know the strategies by which social insects evolved in feature-rich landscapes develop their familiarity with their foraging landscape and whether such spatial familiarity contributes to their future foraging abilities. We postulate that they may invest much effort (for instance, spending greater amount of time out of their nest) during their initial foraging phase to obtain adequate spatial familiarity and spatial memory of rewarding patches. This may eventually enable them to find rewarding sites with much less or even no large-scale searching.

*Ropalidia marginata* is an aseasonal predatory social wasp inhabiting feature-rich tropical landscapes. Females typically survive for 9 to 329 days (mean ± s.d. = 135.9 ± 86.3) in laboratory conditions (Sen and Gadagkar, 2010) and colonies typically consist of 1 to 200 females (mean ± s.d. = 21.9 ± 22.3) (Gadagkar, 2001). Upon eclosion, workers first perform intranidal tasks and then gradually increase the proportion of extranidal tasks (i.e. foraging) with the advancement of their age (Naug and Gadagkar, 1998). They forage solitarily for food (spiders, larvae of other insects etc.), plant fibres (as building materials) and water. If a prey is too big to carry, they cut the prey into pieces and bring those pieces to their nest in multiple bouts. They typically forage within about 500 m from their nests (Mandal and Gadagkar, 2015) and experienced foragers possess comprehensive spatial familiarity with their foraging landscape (Mandal et al., 2017). Each forager has to acquire this spatial familiarity as they forage solitarily. This makes all the foragers of a colony equally suitable for our experiment.

To test whether they spend a substantial amount of time outside their nest during the initial phase of their foraging life (probably for exploring and learning the features of the foraging landscape), and whether they reduce their foraging effort without affecting their foraging benefits at the later stage, we first checked whether the number of trips per day, proportion of time that the forager wasps spent daily on foraging or the average foraging duration per trip change with their age following a pattern that has an initial ascent followed by a descent, and whether these patterns explain the data better than a linear function. In parallel, we checked whether their foraging benefits increased with the advancement of their age. We also predicted that individual foraging wasps, like ants (Wehner et al., 2004) and bees (Capaldi et al., 2000), might develop fidelity for a particular (rewarding) direction to go for foraging with age; young wasps may not develop such a choice for any particular direction during their explorative phase until they come across a rewarding patch (for instance, a food source). We tested this prediction by analysing all the outbound flight directions of individual wasps. Also, though insects rely heavily on path integration during early foraging bouts, with the advancement of their age/experience, they tend to rely more on learnt information acquired from their foraging landscape, if available (Bühlmann et al., 2011; Cheng et al., 2012; Kohler and Wehner, 2005; Menzel and Greggers, 2015; Müller and Wehner, 2010; Narendra, 2007; Wystrach et al., 2014). Since path integration is an error-prone system and the error increases with directional changes during a trip, young and inexperienced foragers are expected to stick to a particular relative direction for a particular trip, i.e. going out and returning along the same path (though they may choose a different direction for subsequent trips) (Capaldi et al., 2000). On the other hand, familiarity with the landscape may enable experienced foragers to explore several places in different directions during a single trip (Hassell and Southwood, 1978). This may be possible by developing preference for habitual foraging directions (Osborne et al., 2013; Woodgate et al., 2016) and eventually using trapline routes (Buatois and Lihoreau, 2016; Lihoreau et al., 2012; Saleh and Chittka, 2007) and novel foraging routes (Menzel et al., 1998). Thus, after each foraging bout, they may return from the direction of sites of their last preference. Therefore, we have checked whether wasps developed any directional preference for their incoming flights and whether the angular difference between the outbound and subsequent inbound directions increased with advancing age.

## MATERIALS AND METHODS

We performed this experiment using three naturally occurring nests (N17, N18 and N21), of *Ropalidia marginata* (Lepeletier 1836) (Vespidae, Polistinae) located at the Indian Institute of Science campus, Bangalore, India (13°01⍰ N, 077°34⍰ E). The landscape of the main campus is spread over an area of about 1.55 km^2^ and dominated by densely distributed trees (with an average height of about 30 m) and shrubs covering about 75% area of the landscape, along with small to medium sized (i.e. maximum height of about 35 m) academic and residential buildings, limiting a continuous view within a maximum distance to about 30 m on ground. Uninterrupted view for much longer distance could only be accessed on roads that had varied lengths with a width of 3-6 m, and with light motor traffic. Such a landscape is of special interest, as foraging wasps typically cruise within a height of 2-10 m from ground and thus can access very small visual catchment area during their regular foraging trips.

We found all the three experimental nests inside electrical-fuse boxes attached to roadside lampposts at a height of about 50 cm from ground. Immediately after locating the nests, we sealed all the holes of those boxes except the one on the frontal lid of the boxes to get the wasps accustomed with it as the only exit and entrance. We also placed a mimic camera set-up 30 cm away from the frontal lid of the boxes to get the foraging wasps used to it and kept it until we replaced it with a real look-alike video camera for data collection (Figure 1). We carried out the experiment in three consecutive steps,

**Figure 1.**
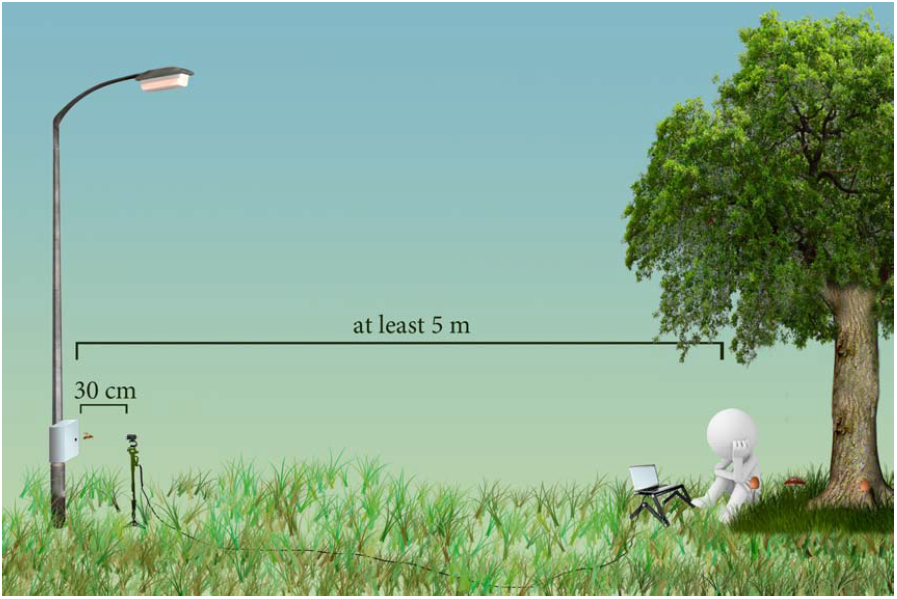
A schematic representation of the experimental set-up. We found all the three experimental colonies of wasps within electric boxes attached to roadside lampposts. By default, these boxes had two holes, one on the frontal lid of the box another at the bottom of the box. We sealed the hole at the bottom so that wasps use the frontal hole as their only entrance and exit. A motion sensitive video camera was placed 30 cm away in front of the frontal hole, so that whenever a wasp came out or went inside the box, the camera started recording the video. The timing of departure and arrival of the wasps, as well as the foraged material could be retrieved from the video. The video was stored into a laptop computer connected to the camera and placed at least 5 m away from the lamppost. An observer, clad in camouflage attire, sat near the laptop and recorded the vanishing direction of the outbound and inbound foraging trips.

### Step 1. Accounting for the age of the wasps

Since we required to know the age of the wasps as the first step of our experiment, after finding a colony, we took daily census of the wasp colonies at night until we knew the age of all the resident wasps of the colony. We uniquely colour-marked all the newly eclosed wasps on their thorax and/or abdomen using Testors^©^ quick drying enamel paints along with recording their date of eclosion. We started taking the daily census of N17, N18 and N21 on October 1, November 11 and December 23, 2013, respectively.

### Step 2. Data collection

Next, we replaced the dummy camera set-up with a real motion sensitive web camera and connected that to a laptop kept at least 5 m away from the lampposts. The camera was set to start recording (with 2 seconds of pre-recording function, using Webcam Zone Trigger ™ software) upon detection of any movement within its field of view (which included the hole on the frontal lid of the box and part of the frontal lid, see Figure 2). Thus, the video captured the identity of all the outgoing/incoming wasps when it appeared within its visual field as well as the time of their departure and arrival, and the identity of foraged materials, if any. An observer sat near the laptop in camouflage attire and recorded data on the vanishing bearing of the outbound and inbound flights performed by the wasps. From such a distance, the observer could see the flight direction of the wasps but not the colour marks on them; hence the observer was practically blind to their identity. We included the vanishing bearings of the outbound wasps in the analysis only if the wasps could be followed for at least 5 m. Similarly, for the inbound wasps, we included data in the analysis only when the observer could first notice the wasp when it was at least 5 m away from their nest. We recorded the vanishing bearings with a 10° interval (by marked transects). For each nest, we collected data for three consecutive days. Each day, we turned on the set-up of video recording at least 5 minutes before sunrise and stopped it after sunset. Thus, we made observation for a total of 30h 45min (10h 15min per day) for N17 on January 29, 30 and 31, 2014, 38h 15min (12h 45min per day) for N18 on April 21, 22 and 23, 2014 and 37h 30min (12h 30min per day) for N21 on April 7, 8 and 9, 2014. The data consisted of 78, 89 and 105 unique foraging wasps from N17, N18 and N21 respectively.

**Figure 2.**
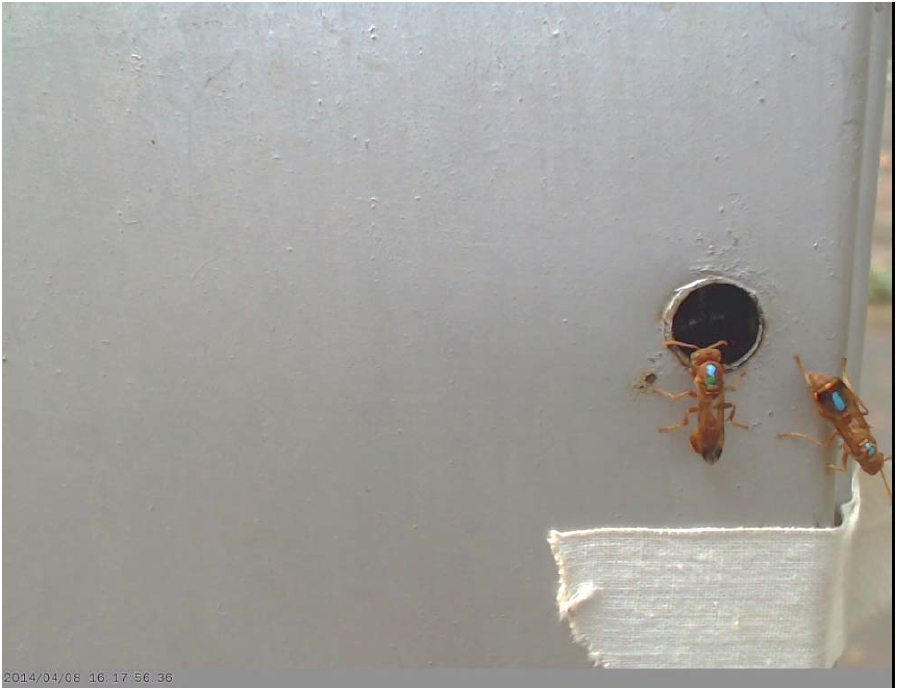
A still frame from the collected video data of the nest N21. In this frame, recorded at 16:17:56.36 hrs (mentioned at the bottom right corner of the frame), a wasp with light blue (coded as L) on top of the thorax and dark green (coded as D) below as well as on her abdomen (hence named as LD) is going inside the box through the exit/entrance hole, while wasp with light blue on thorax as well as on abdomen (hence named as LL) is going out for a foraging trip. None of them is carrying anything.

### Step 3. Data analysis

We performed all statistical analyses using RStudio interface 0.99.891 for R version 3.2.2. For each wasp, we calculated the number of foraging trips she made, proportion of time she spent outside her nest, her average trip duration, foraging benefits (as foraging success and foraging efficiency, see below for definition), consistency in the direction of outbound and inbound trips, and for each trip, the angular difference between the direction of outbound flight and subsequent inbound flight. Next, we checked for any relationships of these parameters with the age of the wasps.

We calculated the proportion of time spent outside the nest as the ratio of the total time that the wasp spent outside her nest during the three days of observation and the total duration of observation on those three days. To check for patterns in the relationships between the proportion of the time spent outside her nest and age of the wasps, we fitted three mathematical functions to the data separately for each of the three nests a linear and a quadratic function with an assumption of symmetry of the data and an asymmetric non-linear rational function namely “Holling type IV” (Bolker,2008). The last was fitted to the data with nonlinear least squares (nls) regression and was expressed as (A×age ^2^)/(B + C×age + age ^2^), where A is the value of the proportion of time spent outside when the curve reaches asymptote, and -2×*B/C* demarcates the age at the peak of the curve. Similarly, we calculated the average duration of foraging trips by dividing the total time a wasp spent on foraging in three days with the total number of trips she had made in these three days and fitted linear, quadratic and Holling type IV function to the data for checking its possible relationships with the age of the wasps.

We computed ‘foraging success’ as ‘the ratio of the number of trips in which a wasp brought food to her nest and the total number of trips she made’. We computed ‘foraging efficiency’ as ‘the number of times the wasp brought food to the nest divided by the proportion of time she spends outside of her nest’. We first fitted a linear and then a quadratic function to explore the relationships between both foraging success and foraging efficiency with the age of the wasps, separately for the three nests.

Next, we tested whether with age, wasps developed a preference toward any particular direction to go out for foraging. We measured their age in days and directional preference as the length of the mean vector (*r* value) (Batschelet, 1981) of all the outbound directions shown by each wasp on a day and averaged over three days. We eliminated those wasps from our analysis for which we had data for only one outbound trip in a day. Thus, our dataset for this analysis comprised of 54 (for N17), 71 (for N18) and 68 (for N21) unique wasps. We fitted a linear and a saturating function namely Michaelis-Menten to the data and checked which one explained the data better. The expression of Michaelis-Menten function was (A × age)/(B + age), where A defines the value of dependent variable (*r* value) at asymptote and B defines the value of the independent variable (age of wasps) at A/2. Similarly, we have checked whether with age, wasps developed any directionality for their inbound trips. Here, the data set comprised of 51, 71 and 80 unique wasps from N17, N18 and N21, respectively. Since we wanted to check whether after an initial increase, wasps reduced the directionality of their inbound trips, we also fitted a quadratic function along with linear and Michaelis-Menten function to the data and checked for the best fit.

To know whether the wasps started taking detours with increasing age, we checked the angular difference between the direction of outbound and subsequent inbound flights for each trip of each wasp. The angular difference ranged from 0° to 180°. Though we agree that reaching to a conclusion about their entire foraging route from the outbound and inbound directional data acquired only from a distance of 5 m around their nests is a bit over-speculation, we still used the data to get a rough proxy of the foraging route during a trip, assuming more angular difference as the indication of greater detour during their foraging trip. Since we had several data points from the same wasp and we had three experimental nests, we fitted a generalized linear mixed effects model (lme4 package) with a Poisson error family and log link, taking data from all the wasps of all the nests together. For better explanatory power of the model, we took the rescaled value (and not transformed value) of the angular difference between outbound and inbound direction as the response variable (using the ‘scales’ package; Wickham, 2016), the age of the wasps as the explanatory variable and the identity of the wasps nested within their colony ID as random effects.

## RESULTS

During the three days of observation, 78, 89 and 105 unique wasps from nest N17, N18 and N21 made a total of 607, 1173 and 2407 foraging trips, respectively. On the basis of the material that a wasp brought back to her nest, most of the wasps (100%, 95.5% and 92.37% from N17, N18 and N21, respectively) could be classified into four categories: wasps that did not bring anything, brought building material, water or food to their nests. However, there were few cases in which a single wasp brought more than one material (see Table S1 in supplementary materials). We found that the water foragers (defined as wasps that brought water more than once and more than anything else) (2, 1 and 8 wasps in N17, N18 and N21, respectively), made much higher numbers of trips than other foragers in fact water foragers were outliers in the number of trips they made. Compared to foragers that brought other things, trip duration of the water foragers was much shorter (GLMM, Estimate: 0.765, z value: 141.72 *P <* 0.01) and their success rate was much higher (χ^2^ test, *P <* 0.01) (Table S1). They showed high directionality for outbound as well as inbound trips (*r* value on any given day was greater than 0.9 for all the water foragers), and had much less angular difference between outbound trips and subsequent inbound trip than the other foragers (GLMM,Estimate: 0.3545, z value: 10.60 *P <* 0.01). Because of these reasons, we excluded water foragers from all the analyses except the analyses that we performed to check for relationships between age of the wasps and the proportion of time they spent outside their nest as well as the average duration of their foraging trips. For an overview of the foraging activities on these three nests, see Figure 3 and Table S1.

**Figure 3.**
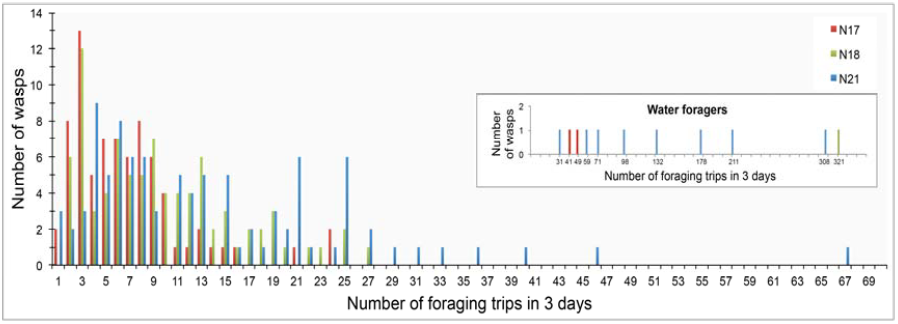
The number of foraging trips performed by wasps from three nests (N17, N18 and N21) during the three days of observation. Number of trips made by the water foragers was much higher than other foragers (see Table S1), and an inclusion of those into this graph would have made visualization of data from other wasps difficult. While data on water foragers are presented in the inset, data on other wasps are in the main graph.

### Change in number of foraging trips per day with age

Wasps from all three nests initially increased and later decreased their number of foraging trips per day with increasing age. A quadratic function explained the rate of change better compared to a linear function (ANOVA, *P <* 0.05) (Figure 4).

**Figure 4.**
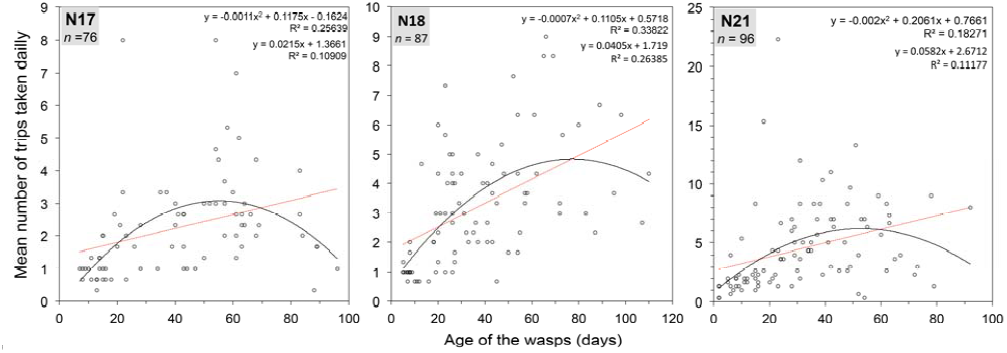
Change in the number of foraging trips taken daily by wasps with their age. For all the three nests, the quadratic model explained more variation of the data than the linear model. Comparison of AIC values of linear (N17=282.36, N18=348.82, N21=512.95) and quadratic model (N17=270.63, N18=341.56, N21=506.96) reconfirms the better fit of the quadratic model (ANOVA, *P <* 0.05). An attempt to fit a Holling type IV function revealed non-significant values of the parameters for all three nests. It is noteworthy that we have not included water foragers in this analysis.

### Change in the proportion of time spent on foraging with age

A linear fit explained only 14.91% (AIC = -64.09), 18.28% (AIC = -8.44) and 26% (AIC = -16.92) of the total variation of the relationship between the age of the wasps and the proportion of time wasps spent outside their nests for N17, N18 and N21, respectively. The quadratic model explained 46.29% (AIC = -98.98), 53.10 % (AIC = - 55.86) and 55.31% (AIC = -67.87) of the total variation of the data from N17, N18 and N21, respectively. When we fitted the function Holling type IV to the data, which has a characteristic of initial rapid ascent and after reaching to a peak, a gradual descent, greater amount of variation in the data was explained with decreased AIC values (N17: 54.46%, AIC = -111.85; N18: 54.74%, AIC = -59.02; N21: 55.98%, AIC = -69.45) (Figure 5). It may be noted that the few young individuals that brought food (red circles in Figure 5) generally spent more time outside of their nest while older individuals that brought food spent varied amount of time outside nest.

**Figure 5.**
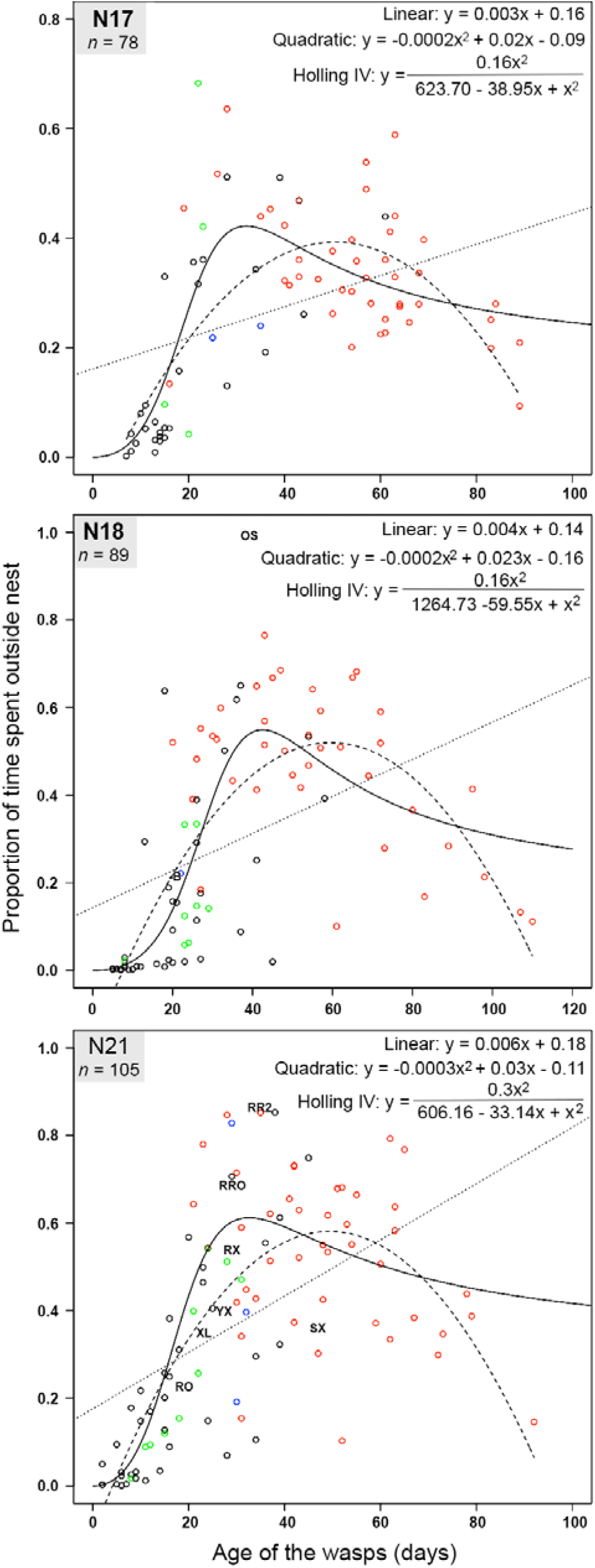
With increasing age, the change in the proportion of the time Ropalidia marginata wasps spent on foraging (time spent outside their nests) in three natural colonies. (namely N17, N18 and N21). In the plots, black circles represent wasps that did not bring anything to their nest during the three days of observation. Likewise, green, blue and red circles represent wasps that brought building material, water and food at least once to their nest, respectively. One wasp (named as „B) in N18 and three wasps (named RO, XL, YX) in N21 brought both building material and water; one wasp (named OS) in N18 brought both water and food; three wasps (RR2, RRO, RX) from N21 brought building material and food; one wasp (SX) from N21 brought building material, water and food. Three mathematical functions (linear, quadratic and Holling type IV) were fitted to the data from each nest. For all the three nests, Holling type IV provided the best fit. For all the parameters of this function, p value was <0.05 for all the nests.

### Change in average foraging duration per trip with age

Similarly, we checked for the relationship between the age and the average foraging duration per trip of the wasps by fitting a linear, quadratic and Holling type IV function to the dataset. Here also, Holling type IV provided the best fit for the data from all three nests; it explained 31.45% (AIC = 857.68) of the variation of the data (compared to 0.124% (AIC =885.04) by linear and 14.51% (AIC = 874.90) by quadratic function) from N17, 41.49% (AIC = 973.52) of the variation of the data (compared to 5.03% (AIC = 1014.62) by linear and 32.4% (AIC = 986.36) by quadratic function) from N18, and 15.45% (AIC = 1164.27) of the variation of the data (compared to 4.08% (1175.52) by linear and 11% (AIC = 1169.66) by quadratic function) from N21 (Figure 6).

**Figure 6.**
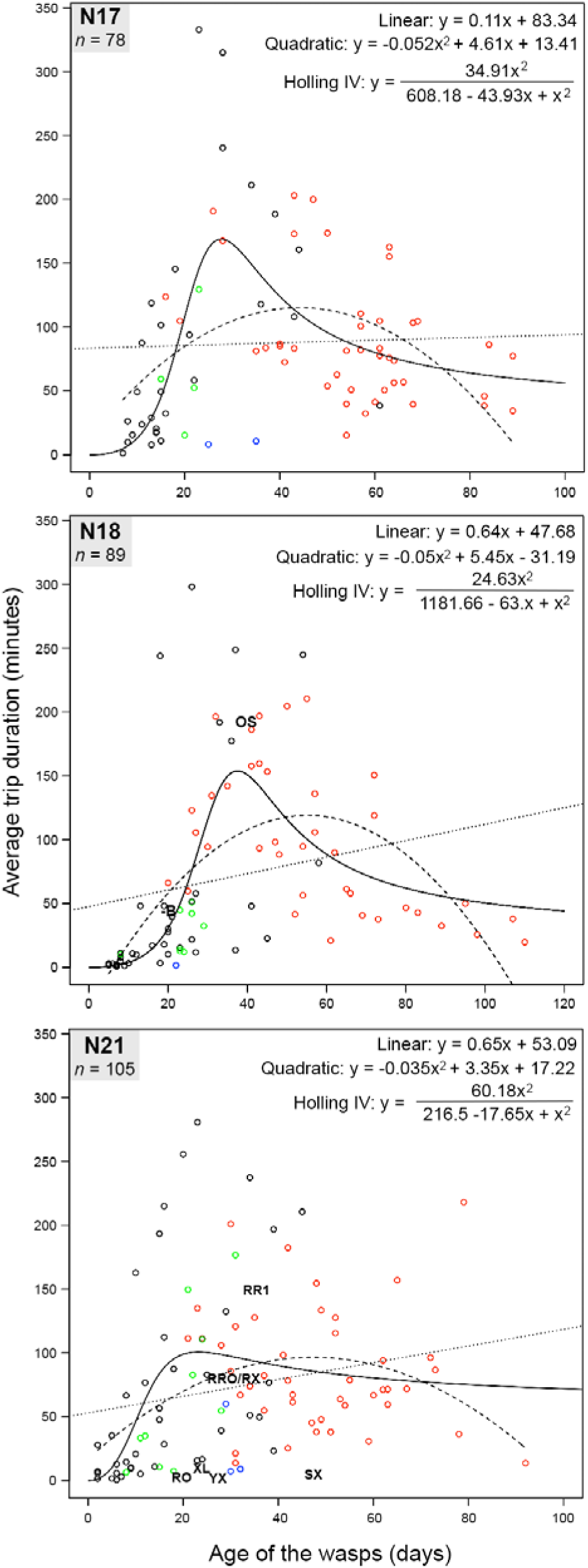
Average foraging duration per trip is also explained best by Holling type IV function, which has a characteristic of rapid initial ascent and after reaching a peak, a gradual descent. For all the parameters of this function, P value was <0.05 for all the nests. As represented in Figure 5, black, green, blue and red circles represent wasps that did not bring anything, and wasps that brought building material, water and food at least once to their nest, respectively during the three days of observation. Similarly, one wasp (named as „B) in N18 and three wasps (named RO, XL, YX) in N21 brought both building material and water; one wasp (named OS) in N18 brought both water and food; three wasps (RR2, RRO, RX) from N21 brought building material and food; one wasp (SX) from N21 brought building material, water and food.

### Change in foraging success with age

With the increase in their age, foraging success (i.e. proportion of trips in which wasps brought food) of wasps from all the three nests increased significantly (*P <* 0.05) (Figure 7). A linear model explained 57.17%, 47.73% and 40.95% of the variation of the data from N17, N18 and N21, respectively. Attempt to fit a quadratic function to the data revealed insignificant *p* values (i.e. *P >* 0.05) for all the parameters for N17 and N21, and a p value less than 0.05 (i.e. 0.027) only for the quadratic term for N18.

**Figure 7.**
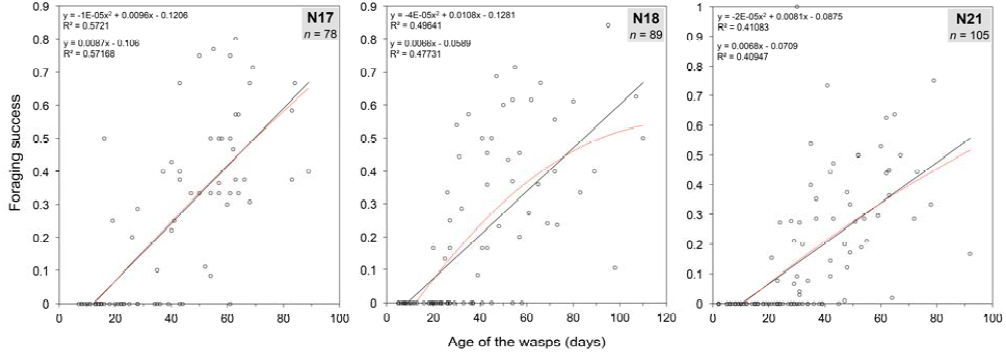
Foraging success (calculated by the ratio of the trips in which a wasp brought food and all the trips that the wasp made) of the wasps increased linearly with increase in their age.

### Change in foraging efficiency with age

Likewise, foraging efficiency (i.e. the number of times a wasp brought food to her nest in unit time she spent on foraging) also increased with the advancement of their age (Figure 8). While all the parameters of linear models for all the nests were significant (*P<* 0.05) (and explained 61.08%, 65.5% and 63% of the variation of the data from N17, N18 and N21, respectively), no parameters of a quadratic mod l were significant for N17, and only the quadratic term was significant for N18 and N21.

**Figure 8.**
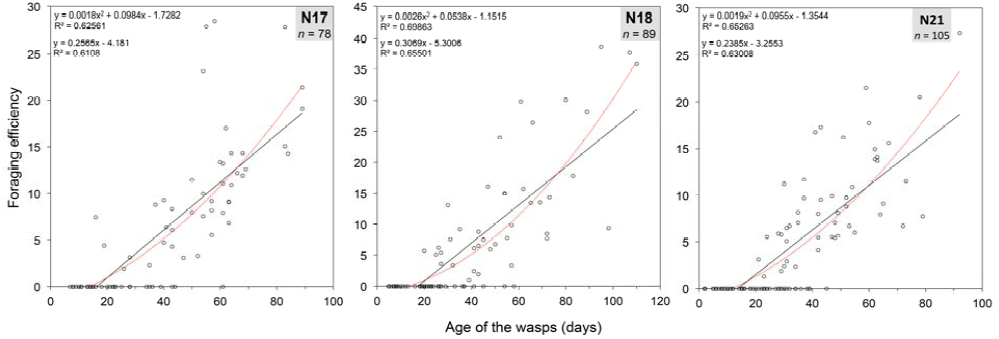
Foraging efficiency, computed by the number of times a wasp brought food to her nest divided by the proportion of time it spent outside of her nest, increased linearly with the advancement of their age. (*P* < 0.05 for all the three nests).

### Developing directional fidelity for outbound trips with age

The average of mean vector length (*r* value) of the outbound flight directions increased with the age of the wasps from all the three nests. We found that Michaelis-Menten function (Figure 9) fitted better than linear function to the data from all the three nests. The former function explained 40.67%, 51.30% and 54.58% of the total variation of the data from N17, N18 and N21 respectively (AIC: -9.17 for N17, -26.66 for N18 and -48.01 for N21) whereas the latter explained 22.9%, 42.55% and 54.74% of the total variation in the data from N17, N18 and N21 respectively (AIC: 4.98 for N17, -15.95 for N18, -33.81 for N21).

**Figure 9.**
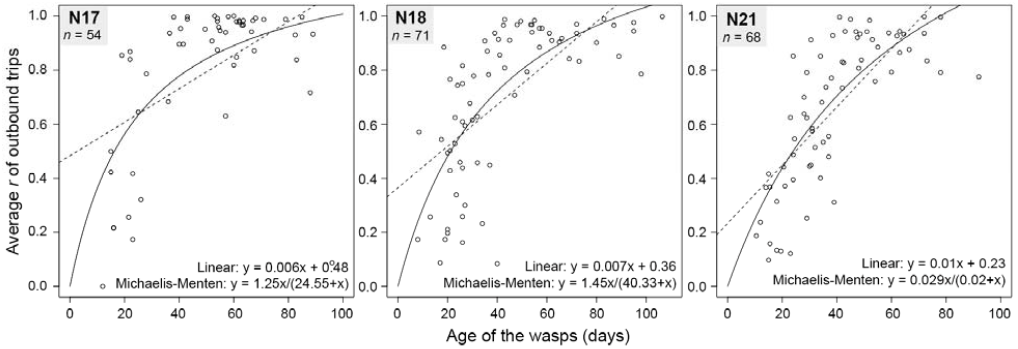
Directional fidelity to go for foraging, calculated by averaging the mean vector length of the outbound directions (*r*) shown by each wasp each day, increased with age of the wasps following a Michaelis-Menten function, average *r* = (A × age)/(B + age). For all the values of A and B for all the nests, *P* < 0.05.

### Developing directional fidelity for inbound trips with age

Interestingly, the average of the mean vector length (*r* value) of the inbound flights made by wasps also increased with their age. Whereas the Michaelis-Menten function best explained the data from N17 and N18, a quadratic function best explained the data from N21 (Figure 10). While the linear function explained 16.92% (AIC=-14.44), 16.64% (AIC=-18.72), 13.91% (AIC=-18.48) of the variation of the data from N17, N18 and N21 respectively, a quadratic function explained 24.6% (AIC=-17.38), 30.91% (AIC=-30.05), 43.19% (AIC=-49.73), and the Michaelis-Menten function explained 24.6% (AIC=-19.36), 32.07% (AIC=-33.25), 37.02% (AIC=-43.48) of the variation of the data from these three nests.

**Figure 10.**
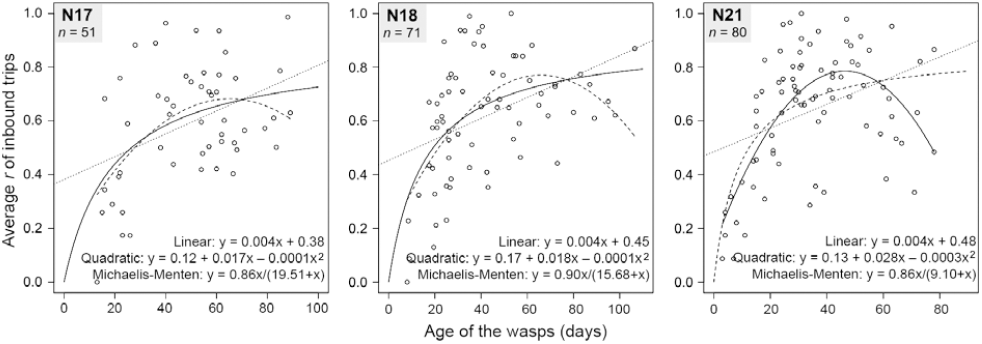
Directional fidelity of the inbound trips increased with age. Though slopes and intercepts of linear functions were significant (*P <* 0.05) for all the three nests, data from N17 and N18 were best explained by the Michaelis-Menten function (*P <* 0.05 for all the parameters), and data from N21 were best explained by a quadratic function (*P <* 0.05 for all the parameters).

### Change in angular difference between outbound and subsequent inbound direction with age

When we checked for the relationship between the angular difference in the outbound and subsequent inbound flight direction with the age of the wasps using a generalized linear mixed model, we found that the angular difference increased with the advancement of age. In the model, we found the estimate of intercept as 3.55 ± 0.11 (SE), and estimate of age as 0.009 ± 0.002 (SE), *P <* 0.05 for both intercept and age.

## DISCUSSION

In a previous study, we have shown that spatial familiarity with their foraging landscape boosts homing performance in the tropical social wasp *Ropalidia marginata* significantly (Mandal et al., 2017). The same has also been shown in several other social insects (Collett et al., 2013; Narendra et al., 2013; Palikij et al., 2012). Possessing spatial familiarity with the foraging landscape can enable animals to minimize foraging effort without affecting their foraging benefits, which can be a great advantage in natural contexts. In an attempt to investigate the ontogeny of spatial familiarity with foraging landscape and foraging capabilities in the tropical social wasp *Ropalidia marginata*, especially to know whether *R. marginata* foragers reduce their foraging effort but increase their foraging gain with an increase in their experience with the foraging landscape, first we checked how the number of foraging trips per day, average duration of their foraging trip and the total time they spent on foraging changed with the advancement of the age of free-foraging wasps from three nests, along with recording the materials that they brought to their nest. We found that most of the wasps started going out of their nest within about the first two weeks of their lives. They began their foraging life with few short duration trips and gradually increased the number of foraging trips per day till about the middle of their foraging career, and then they reduced that. The relationship between the number of trips per day and their age was best explained by a quadratic function. During their initial foraging life, they also rapidly increased the average duration of their foraging trips and the total time that they spent daily on foraging. However, after about four weeks of age, their average foraging duration per trip and the proportion of time they spent in foraging per day followed a gradual decrease. Such an initial rapid increase followed by a gradual decrease in their foraging effort indicate that wasps invest great amount of effort in the initial phase of their foraging life probably for learning/exploring their foraging landscape; once the foragers acquire sufficient familiarity with their foraging landscape, they decrease their foraging effort.

It is noteworthy that during this initial phase of learning/exploring, which is till about four weeks of their age, they rarely brought food to their nests although several wasps brought building materials and water. Compared to desert ants (Narendra et al., 2013; Wehner et al., 2004), honey bees (Capaldi and Dyer, 1999) and bumblebees (Osborne et al., 2013; Woodgate et al., 2016), which start bringing food to their nest after only a few exploration flights/walks, the exploration period in this wasp seems to be much longer. However, this is not surprising if the duration of the exploratory phase can be contributed to the complexity of the foraging landscape and the distribution of the foraged materials. Compared to tropical insects like this wasp, desert ants, for instance, encounter fewer visual features to learn from their foraging landscape (which include occasional small bushes and minor variation on the surface of the land) and so once they get accustomed to being outside their nest, they get ready to forage for food depending heavily on path integration system. The chance of encountering food, i.e. dead insects may also be higher for these short-lived thermophilic ants in such an extreme environment than finding camouflaging prey by the wasps in tropics. Thus, these ants can start foraging for food much earlier.

On the other hand, *R. marginata* wasps live for much longer time and typically bring building materials or water before bringing food to their nest (Naug and Gadagkar, 1998). Such age-based polyethism has been reported in other paper-wasps as well (Jeanne and Taylor, 2009). Apparently building materials (i.e. plant fibres) were abundantly available in all the directions throughout the landscape. Thus, to collect building materials, wasps need not to go for any particular place and foragers with very little experience of the landscape could also accomplish this task. In fact, we found that the foragers that collected building materials were indeed among the youngest foragers of their colony. Compared to finding building material, finding a source of water needed more exploration and bringing water repeatedly from the same place implies that they have learnt the location of the water source. We found that the water foragers had several successful short-duration trips per day and the mean direction of their outbound flights was toward directions in which there was at least one permanent source of water (for instance, fountain or small water reservoir) within about 100 m. Wasps that brought food were among the oldest individuals of their colony. This may be because finding a prey may need much more searching, which includes visits to several places with potential of prey-availability. Several times, wasps took great time (more than 2 hours) to bring food once, followed by few trips when they brought food quite quickly (about 10 minutes). Several times, they brought food spending more or less equal amount of time on all the trips (all more than 2 hours). When they take greater time to bring food to their nest, they probably first find the prey by searching and then hunt it. Next time onwards, they bring the pieces of the kill, so it takes much less time. Also, since their kill is lucrative food for several ants, wasps might hunt for bigger prey when they are close to their nests (so that they can reach back quickly to the hunting site before ants claim the kill) but may kill only smaller prey (which can be carried in a single bout) when they are far from their nest. Thus, to collect food, a thorough familiarity with the foraging landscape might be very necessary for the wasps.

The European honey bees and bumblebees inhabit temperate landscapes that are more complex than deserts but less complex than tropics in which the wasps inhabit, and forage for nectar/pollen. Additionally, flowers are stationary and advertise themselves to attract pollinators like bees, contrary to the mobile prey of the wasps that probably attempt to not getting discovered by the wasps. Also, unlike these wasps that practice solitary foraging, honey bee foragers can get information about rewarding patches from a nest-mate. Putting all together, wasps might need to search for food much more vigorously than bees need to do - bees may not need to acquire a detailed familiarity of their foraging landscape to find their food and they may acquire the required familiarity with the foraging landscape within much shorter time enabling them to start bringing food early in life. Thus, a prolonged exploratory phase can be expected for a predatory wasp that inhabits highly dense tropical landscape and does not have the advantage of conspecific recruitment. However, such a high investment in learning during early foraging life can only be balanced if the forager wasps (can use the acquired spatial familiarity to) increase their foraging benefits with their increasing age.

Interestingly, we found that both foraging success (measured by the ratio of the number of trips in which a wasp brought food and all the trips that she made) and foraging efficiency (measured by the number of times a wasp brought food per unit time she spent on foraging) of the wasps increased with their age. The reduction in their foraging effort and the increase in foraging benefit with increasing age indicate learning and memorising of the features of the landscape in early foraging life, and using the spatial familiarity later for foraging; this might be a stable strategy for predatory animals inhabiting highly dense landscapes. On the contrary, although desert ants (Wehner et al., 2004) and honey bees (Dukas and Visscher, 1994) are also known to increase their foraging benefits with the advancement of their foraging life, both of these insects do so by increasing their foraging effort; desert ants increase their number of foraging trips (Wehner et al., 2004) and honey bees increase their foraging speed and distance (from hive) with increasing age (Capaldi et al., 2000).

In a landscape where food is randomly distributed into patches, animals can achieve foraging competence by memorising the locations of rewarding patches and reaching those places directly (instead of searching for prey every time randomly). We found that wasps indeed developed directional fidelity for the outbound flights with increasing age. The relationship between the consistency in their outbound direction and their age can be explained best by the saturating Michaelis-Menten function. This suggests that the wasps fly in many directions, probably for learning/exploring in the initial foraging stages (and therefore show less directional fidelity), but soon develop a choice for particular direction, maybe after encountering prey in that particular direction. Wasps prefer to begin their search for prey in that direction or in a direction we they have encountered pray in recent past, at least for some days. The saturation of high directional preference in the older wasps may also indicate that a rewarding patch continues to be rewarding at least for three days.

Since young wasps do not show any directional preference for their outbound trips, if they return from the same direction, they should not have any directionality for their inbound flights as well, and the angular difference between the outbound and subsequent inbound flights should be low. On the other hand, since experienced foragers develop a directional preference for outbound trips, if they forage by following a trapline foraging route as bees are known to do (Buatois and Lihoreau, 2016; Lihoreau et al., 2012; Saleh and Chittka, 2007), they should also show an increase in the directionality of inbound trips. However, following more than a single trapline may again reduce the directionality of the inbound trips. With age, wasps from two nests in fact showed an increase in their directionality of inbound trips following the saturating Michaelis-Menten function and wasps of the other nest showed an initial increase followed by decrease in older age. Though our data on the direction of the inbound flights were collected on the basis of the very last stages of the flights and therefore may just indicate the best approach to the nest (and might be unrelated to foraging path), these results might also suggest that depending on the distribution of resources, which in turn depends on many factors including the landscape, wasps might develop one or more trapline foraging routes in different directions. Quite interestingly, we have also found that the angular difference between the direction of outbound flight and subsequent inbound flight increased with the advancement of age of the wasps. This suggests that the young foragers returned from the same direction in which they had their outbound foraging trip, whereas the older wasps returned to their nest from directions different from the direction of their outbound trip. This may have happened because during their learning phase, young wasps probably relied mostly on error-prone path integration, the degree of error of which increases with the change of the angular direction in a single foraging trip. Thus, young foragers did not make much angular deviation during a single trip; similar results has been shown in honey bees (Capaldi et al., 2000). With experience, wasps may enhance their skill in path integration and also they may start using the acquired familiarity with the foraging landscape much more efficiently for navigational guidance, as observed in ants (Mangan and Webb, 2012) and honey bees (Zhang et al., 1999).

## CONCLUSIONS

All these results indicate that these wasps acquire spatial familiarity with their foraging landscape in their initial phase of foraging. They do so probably by learning and developing a long-term memory of the features of the landscape. This memory in turn enables the wasps to reduce their foraging effort and yet increase their foraging benefits. Such an ontogeny of foraging capability that is strongly influenced by the memory of their surrounding landscapes may be a reflection of their evolution in the landmark-rich tropical ecosystem.

## Acknowledgements

We thank Kavita Isvaran for her help with statistical analyses, and Thomas S. Collett for many helpful discussions.

## Disclaimer

SM and RG designed the study, SM and AB conducted the study, SM analysed the data and SM and RG co-wrote the paper. The authors declare no competing interest. The research adheres to the guidelines for the Use of Animals in Research, the legal requirements of the country in which the work was carried out, and all institutional guidelines.

---

**Supplementary materials: Table S1.**
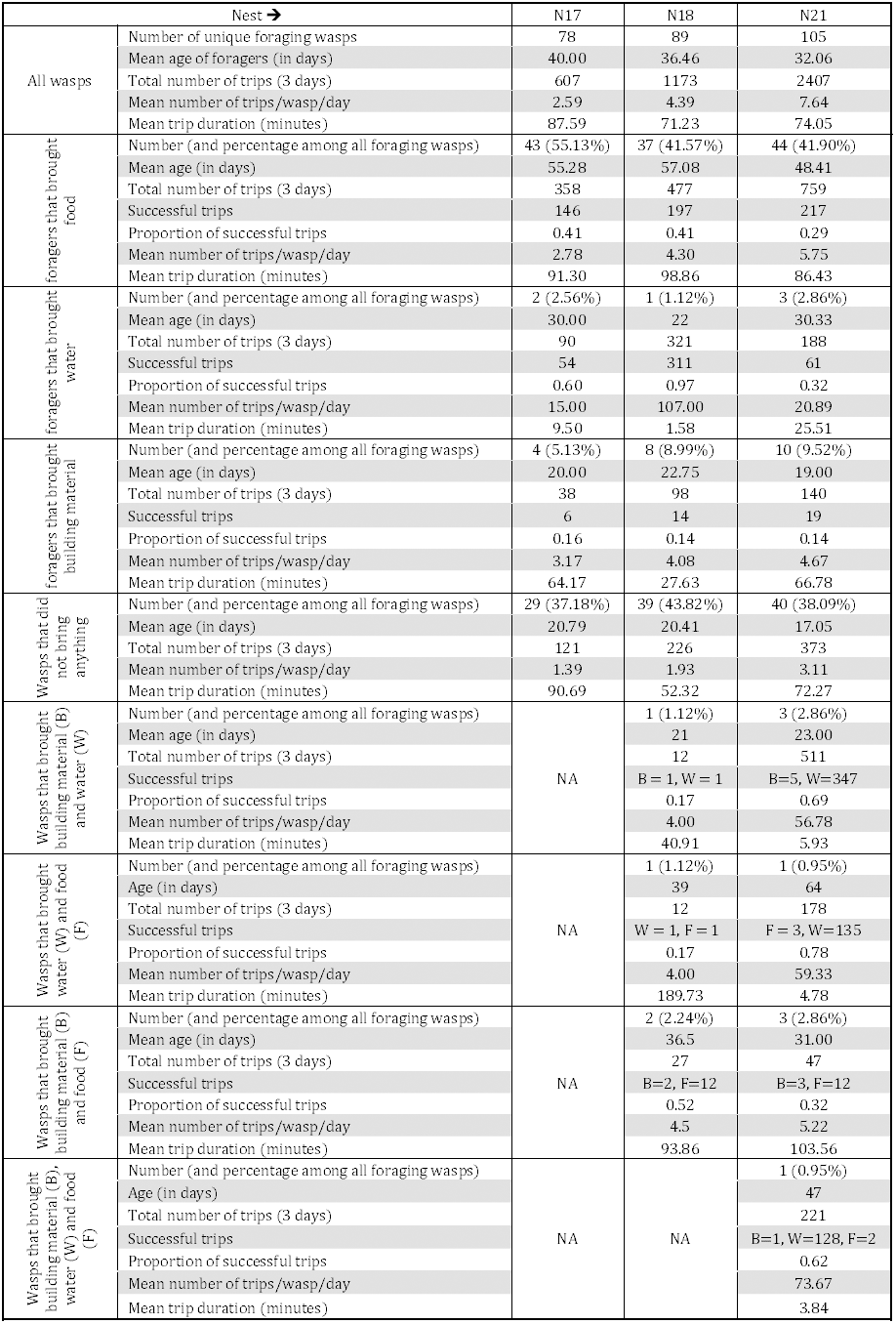
Overview of the data collected from three wasp nests.

